# Dopamine D2R upregulation in nucleus accumbens indirect pathway does not affect Pavlovian or Go/No-Go Learning

**DOI:** 10.1101/2020.04.28.066670

**Authors:** Kelly Martyniuk, Michelle Dandeneau, Peter Balsam, Christoph Kellendonk

**Affiliations:** Department of Neuroscience, Graduate School of Arts and Sciences, Columbia University; Department of Psychiatry, Columbia University; Department of Pharmacology, Columbia University; Division of Molecular Therapeutics, New York State Psychiatric Institute; Division of Developmental Neuroscience, New York State Psychiatric Institute; Department of Psychology, Barnard College

## Abstract

Ventral striatal dopamine is thought to be important for associative learning. Dopamine exerts its role via activation of dopamine D1 and D2 receptors in the ventral striatum. Upregulation of dopamine D2R in the indirect pathway of the nucleus accumbens (NAc) impairs incentive motivation via inhibiting synaptic transmission to the ventral pallidum. Here, we determined whether upregulation of D2Rs and the resulting impairment in indirect pathway function modulates associative learning in an auditory Pavlovian reward learning task as well as Go/No-Go learning in an operant based reward driven Go/No-Go task. We found that upregulation of D2Rs in indirect pathway neurons of the NAc did not affect Pavlovian learning or the extinction of Pavlovian responses, and neither did it alter No-Go learning. A delay in the Go component of the task however could indicate a deficit in learning though it may be attributed to locomotor hyperactivity of the mice. In combination with previously published findings our data suggest that D2Rs in the NAc core play a specific role in regulating motivation by balancing cost/benefit computations but do not necessarily affect associative learning.

## Introduction

The role of ventral striatal dopamine and its receptors in the regulation motivation and learning has been an intensive area of study for the last decades. Pharmacological studies have uncovered an important role for dopamine receptors in the Nucleus accumbens (NAc) in the regulation of incentive motivation and the willingness to work for reward (Aberman, Ward, & Salamone, 1998; Berridge, 2007; Salamone, Correa, Farrar, & Mingote, 2007). In this context dopamine is thought to regulate effort-related processes that are important to overcome work-related response costs rather than to adapt the animals response to changes in reward value (Filla et al., 2018; Hamid et al., 2016; Kelley, Baldo, Pratt, & Will, 2005; Ostlund, Wassum, Murphy, Balleine, & Maidment, 2011; Phillips, Walton, & Jhou, 2007; Salamone et al., 2007; Wanat, Kuhnen, & Phillips, 2010). Upregulation of dopamine D2 receptors (D2Rs) in the adult NAc core enhances performance in a progressive ratio and a concurrent choice tasks that probe for incentive motivation and effort related decision making (Donthamsetti et al., 2018; Gallo et al., 2018; Trifilieff et al., 2013). Notably, cell specific upregulation of D2Rs in indirect pathway (iSPNs) projection neurons (D2R-OE_NAcInd_ mice) is sufficient to enhance motivation, whereas upregulation in cholinergic interneurons (D2R-OE_ChAT_ mice), which also express D2Rs had no effect on progressive ratio performance (Gallo et al., 2018) (Note, that for simplicity we are using the term indirect pathway, a term that has recently been challenged for the NAc (Kupchik et al., 2015).

D2Rs in iSPNs are transported to axonal terminals, where they reduce inhibitory transmission at intra-striatal collaterals and striato-pallidal synapses (Cooper & Stanford, 2001; Dobbs et al., 2016; Floran, Floran, Sierra, & Aceves, 1997; Kohnomi, Koshikawa, & Kobayashi, 2012; Tecuapetla, Koos, Tepper, Kabbani, & Yeckel, 2009). Slice physiological recordings revealed that D2R upregulation in iSPNs enhances this modulation by dopamine. Thus, D2R-OE_NAcInd_ mice display decreased baseline synaptic transmission and an enhanced inhibition of synaptic transmission by D2R activation (Gallo et al., 2018). As you would expect this effect was recorded at intra-striatal collaterals to the direct pathway and the canonical projections to the ventral pallidum (Gallo et al., 2018). A follow up *in vivo* physiological analysis showed that the effects of disinhibition in D2R-OE_NAcInd_ mice are mostly measurable at the level of the striato-pallidal synapse (Gallo et al., 2018). Furthermore, selective inhibition of iSPN synapses in the ventral pallidum is sufficient to enhance progressive ratio performance suggesting that NAc D2Rs promote incentive motivation via enhanced inhibition of striato-pallidal transmission (Gallo et al., 2018).

Ventral striatal dopamine has also been implicated in associative learning. Dopamine neurons have been shown to encode a reward prediction error providing a teaching signal that is required for learning and that is thought to be transmitted to the NAc via the release of dopamine (Day, Roitman, Wightman, & Carelli, 2007; Schultz, Dayan, & Montague, 1997; Steinberg et al., 2013). D2R-OE_NAcInd_ mice should be more sensitive to this signal so that dopamine released in response to a reward predicting cue leads to a stronger inhibition of synaptic transmission, which could affect associative learning. To address this hypothesis, we tested D2R-OE_NAcInd_ mice in Pavlovian conditioning, an associative learning task. In this task mice learn that an auditory stimulus (conditioned stimulus: CS+) predicts the delivery of a food reward, whereas a different auditory stimulus (CS-) is not reinforced. Importantly, CS+ presentation leads to the release of dopamine when mice are acquiring the task (M. R. Bailey et al., 2018). We then extinguished the importance of the CS+ by adding 5 days of extinction training in which animals were not rewarded.

As inhibition of the indirect pathway has been shown to increase response initiation, we further hypothesized that D2R upregulation impairs learning if actions must be suppressed (Carvalho Poyraz et al., 2016). We thus tested D2R-OE_NAcInd_ mice in an instrumental Go/No-Go learning task where in a first step mice learn to press a lever in the presence of a visual stimulus. In a second step they then must learn to withhold from pressing the lever when the stimulus is absent. Last, we measured the activity of D2R-OE_NAcInd_ mice in an open field to determine the functionality of the upregulated receptors in these new cohorts of mice.

We replicated previous findings showing hyperactivity in the open field (Donthamsetti et al., 2018; Gallo et al., 2015). In contrast to our expectations, D2R upregulation neither affected Pavlovian, extinction nor Go/No-Go learning suggesting that D2R upregulation in NAc iMSN enhance motivation (Donthamsetti et al., 2018; Gallo et al., 2015) but does not affect Pavlovian or No-Go learning.

## Methods

### Animals

Adult male and female *Drd2-*Cre BAC transgenic mice (ER44; GENSAT) backcrossed onto the C57BL/6J background were group housed under 12-h light/dark cycle. All experimental procedures were conducted following NIH guidelines and were approved by Institutional Animal Care and Use Committees by Columbia University and New York State Psychiatric Institute. We chose *Drd2-*Cre over A2A-Cre mice to recapitulate the conditions in which we saw enhanced progressive ratio performance. Also, Cre levels are higher in *Drd2-*Cre mice as they can be visualized with anti-Cre immunohistochemistry (Cazorla et al., 2014; Gallo et al., 2018), whereas we cannot detect Cre expression in A2A-Cre mice using the same anti-serum. In our hands *Drd2-Cre* mice leads to recombination of AAV expression constructs in about 5% to 10% of ChAT neurons (Carvalho Poyraz et al., 2016; Gallo et al., 2018).

### Stereotaxic Surgery

Mice (≥8 weeks old) were bilaterally injected with a previously characterized Cre-dependent double-inverted open reading frame (DIO) adeno-associated viruses (AAVs) encoding D2R-ires-Venus, EGFP, (UNC Vector Core, Chapel Hill, NC) into the nucleus accumbens (NAc) using stereotactic Bregma-based coordinates: AP, + 1.70 mm; ML, ± 1.20 mm; DV, –4.1 mm (from dura). Groups of mice used for experiments were first assigned their AAV-genotype in a counterbalanced fashion that accounted for sex, age, home cage origin.

### Operant Apparatus

Eight operant chambers (model Env-307w; Med-Associates, St. Albans, VT) equipped with liquid dippers were used. Each chamber was in a light- and sound-attenuating cabinet equipped with an exhaust fan, which provided 72-dB background white noise in the chamber. The dimensions of the experimental chamber interior were 22 × 18 × 13 cm, with flooring consisting of metal rods placed 0.87 cm apart. A feeder trough was centered on one wall of the chamber. An infrared photocell detector was used to record head entries into the trough. Raising of the dipper inside the trough delivered a drop of evaporated milk reward. A retractable lever was mounted on the same wall as the feeder trough, 5 cm away. A house light located on wall opposite to trough illuminated the chamber throughout all sessions.

### Dipper Training

Four weeks after AAV surgery, mice underwent operant training. Mice were weighed daily and food-restricted to 85–90% of baseline weight; water was available ad libitum. In the first training session, 20 dipper presentations were separated by a variable inter-trial interval (ITI) and ended after 20 rewards were earned or after 30 min had elapsed, whichever occurred first. Criterion consisted of the mouse making head entries during 20 dipper presentations in one session. In the second training session, criterion was achieved when mice made head entries during 30 of 30 dipper presentations.

### Pavlovian Conditioning

Mice were trained for 16 consecutive days in a Pavlovian conditioning paradigm, which consisted of 12 conditioned stimulus-positive (CS+) trials and 12 CS-trials occurring in a pseudorandom order. Each trial consisted of an 80-dB auditory cue presentation for 10 sec, of either a 3 kHz (Cohort 1) or 8 kHz (Cohort 2) tone or white noise (counterbalanced between mice) and after cue offset a milk reward was delivered only in CS+ trials, whereas no reward was delivered in CS-trials. There was a 100 sec variable intertrial interval, drawn from an exponential distribution of times. Head entries in the food port were recorded throughout the session, and anticipatory head entries during the presentation of the cue were considered the conditioned response. The differential score (Head entries/sec (CS – (ITI))) was calculated using either the 2 sec or 10 sec of ITI proceeding the cue. We used 10 sec of ITI to calculate the differential score during the entire 10 sec cue (**Figure 2b**) and 2 sec ITI for the 2 sec binning **(Figure 2d & d**).

**Figure 1.**
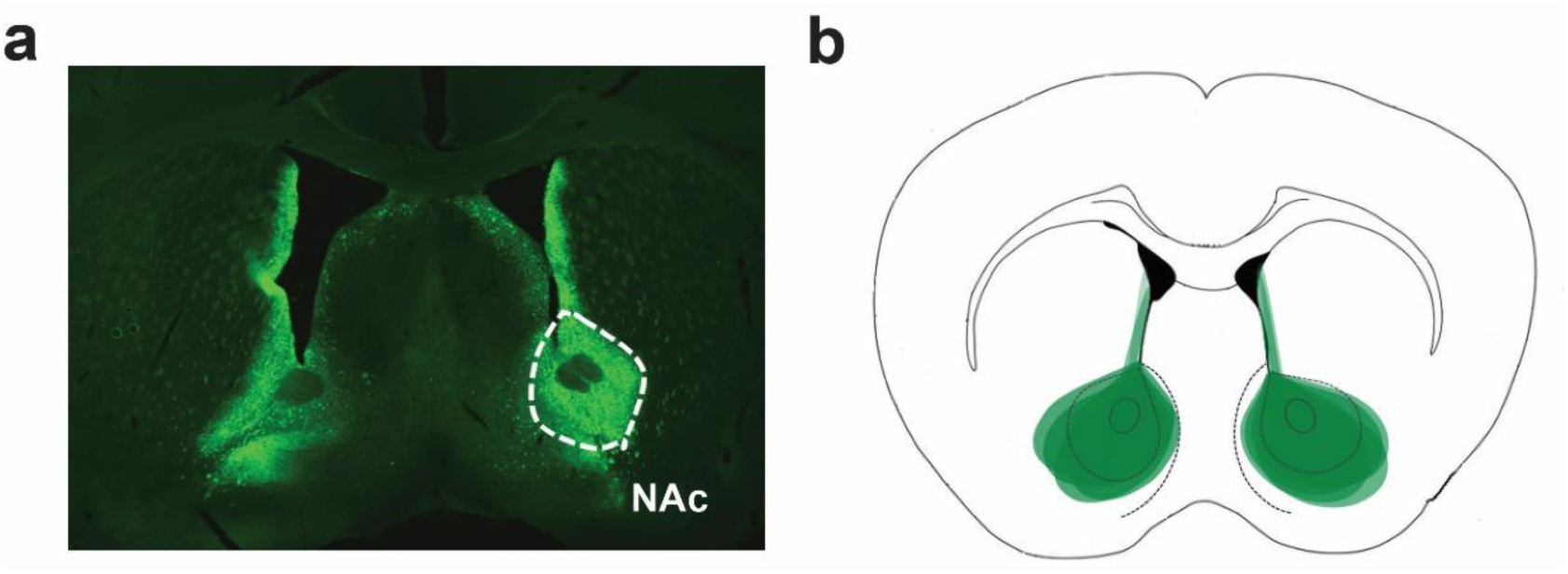
Confirmation of viral spread. **(a)** Coronal section showing D2R-mVenus expression in the NAc of a *Drd2*-Cre mouse. **(b)** Superimposed traces of viral spread from coronal sections at ∼1.0 mm anterior to bregma for all D2ROE_NAcInd_ mice injected with the AVV1-hSyn-DIO-D2R-mVenus into the NAc.

**Figure 2.**
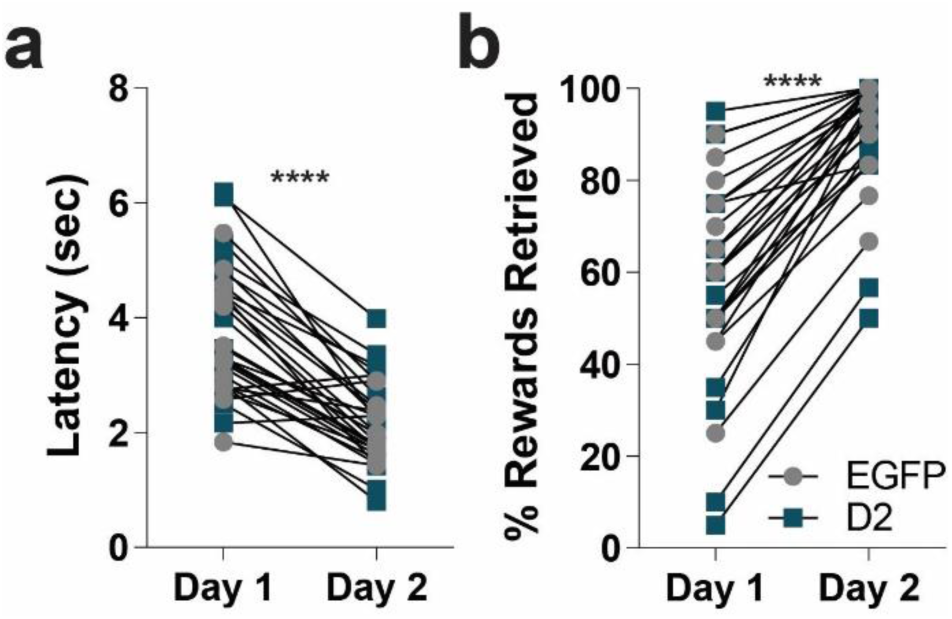
D2R upregulation does not impair trough training. **(a)** Latency to retrieve an unexpected milk reward via head entry into a retractable dipper significantly decreased over 2 days of training for both D2R-OE_NAcInd_ (dark squares) and EGFP_NAcInd_ (light circles) mice (**** *p*< 0.0001). **(b)** Total number of rewards retrieved significantly increased from Day 1 to Day 2 for both groups (**** *p*< 0.0001). Data from 16 animals per genotype was used to calculate all statistics reported.

### Continuous Reinforcement schedule (CRF)

For lever press training, lever presses were reinforced on a continuous reinforcement (CRF) schedule. Levers were retracted after each reinforcer and were presented again after a variable ITI (average 40 sec). The reward consisted of raising the dipper for 5 sec. The session ended when the mouse earned 60 reinforcements, or one hour elapsed, whichever occurred first. Sessions were repeated daily until mice achieved 60 reinforcements.

### Go/No- Go schedule

Mice were first trained on Go trials in which they were required to press a lever within 5 sec of its presentation to receive a reward. If the 5 sec elapsed with no response, the lever would retract, no reward would be presented, and a new ITI (average 40 sec) would begin. Mice were trained on these 5 sec Go-only trials until they reached 75% accuracy over three consecutive days. Once this criterion was achieved, No-Go trials were added in which the lever was presented simultaneously with two cues (the house lights turning off, and a small LED light above the lever turning on). A lack of any lever press within 5 sec, resulted in a reward. A lever press during this period caused the lever to retract, the house lights to turn on, the LED light to turn off, and a new ITI to begin without any reward for that trial. In each session, 30 Go trials were interspersed with 30 No-Go trials presented pseudo-randomly such that there were an equal number of both kinds of trials in every block of 10 trials. Mice were tested for 35 days, and false alarm rate and hit rate were analyzed.

### Locomotor Activity

*D2*-Cre mice injected with D2R- or EGFP-expressing AAVs were tested in open field boxes equipped with infrared photobeams to measure locomotor activity (Med Associates, St. Albans, VT). Data were acquired using Kinder Scientific Motor Monitor software (Poway, CA) and expressed as total distance traveled (cm) over 90 min.

### Histology

Mice were transcardially perfused with ice-cold 4% paraformaldehyde (Sigma, St. Louis, MO) in PBS under deep anesthesia. Brains were harvested, post-fixed overnight and washed in PBS. Free-floating 50-µm coronal sections were obtained using a Leica VT2000 vibratome (Richmond, VA). After incubation in blocking solution (5% horse serum, 0.5% bovine serum albumin in 0.5% PBS-Triton X-100) for 2 hr at room temperature, sections were labeled overnight at 4 °C with primary antibodies against GFP (chicken; 1:1000; AB13970 Abcam, Cambridge, MA). Sections were incubated with fluorescent secondary antibodies (Goat anti-chicken, A488, A11039, ThermoFisher) for 2 hr at RT. Sections were then mounted on slides and cover slipped with Vectashield containing DAPI (Vector, Burlingame, CA). Digital images were acquired using a Zeiss epifluorescence microscope.

### Statistical analysis

Data are expressed as mean ± SEM. Students’ *t*-tests were used to compare between two groups. Multiple comparisons were evaluated by one-way, two-way, or three-way repeated measures ANOVA, using GraphPad Prism software. Statistical significance was considered for *p* <0.05.

## Results

To test the effects of D2R upregulation in iSPNs on Pavlovian Conditioning and Go-No Go learning we generated two cohorts of mice. Cohort 1 was first tested in the Pavlovian Conditioning task followed by the Go/No-Go task. Cohort 2 was first tested in the Pavlovian Conditioning task followed by an extinction procedure. At the end of behavioral testing both cohorts were run in the open field as a positive control. Adult D2R-OE_NAcInd_ mice were generated by injecting a Cre-depending AAV1 expressing D2R-ires-mVenus or GFP into the NAc core of Drd2-Cre mice that express Cre recombinase in iSPNs. This leads to a 3-fold increase in D2 receptor levels in the NAc (Gallo et al., 2018). **Figure 1a** shows a representative image of a coronal section from a mouse with bilateral expression of the D2R-ires-mVenus virus in the NAc. **Figure 1b** shows the spread of virus-mediated expression. We see dense viral expression in the NAc core with some leakage into the lateral NAc shell and dorsal medial striatum (DMS).

Four weeks after AAV injections mice were tested in the Pavlovian conditioning. First, mice were trained in an operant box to retrieve a food reward (evaporated milk) from an automatic dipper. Over the two training days, both groups learned that the dipper provided a reward (**Figure 2**). While there was no difference between the two groups, EGFP_NAcInd_ and D2R-OE_NAcInd_ mice decreased their latency to retrieve the milk reward (**Figure 2a**, *t*= 7.302, *p*<0.0001 and *t=* 6.525, *p*<0.0001, respectively) and increased the total number of rewards retrieved (**Figure 2b**, p<0.0001 Wilcoxon Rank Sum and p<0.0001 Wilcoxon Rank Sum, respectively). These results indicate that D2R upregulation in iSPNs does not affect reward retrieval.

During auditory Pavlovian conditioning mice were presented with two 10 second auditory cues (tone versus white noise; counterbalanced between viral groups). After cue offset of the conditioned stimulus (CS+), a milk reward was delivered whereas no reward was delivered after the offset of the unconditioned stimulus (CS-). **Figure 3a** provides a schematic of the task. Both groups of mice increased their head entries during the CS+ presentation over the 16 days of training (**Figure 3b**). In contrast, during the CS-, head entries increased slightly during the first 4 days and then went down (**Figure 3b**). Both groups were able to distinguish between the CS+ and CS- (RM ANOVA: F(1,15)= 341.6, *p*< 0.0001) and demonstrated learning across 16 days of training (RM ANOVA: F(1,15)= 4.744, *p* <0.0001). However, there were no significant difference in the rate of anticipatory head poking during the CS+ or CS- between D2R and EGFP expressing mice over the 16 days of training (**Figure 3b**, RM 3-way ANOVA: F(15,15)= 0.2672, *p*= 0.9977, N=16/group).

**Figure 3.**
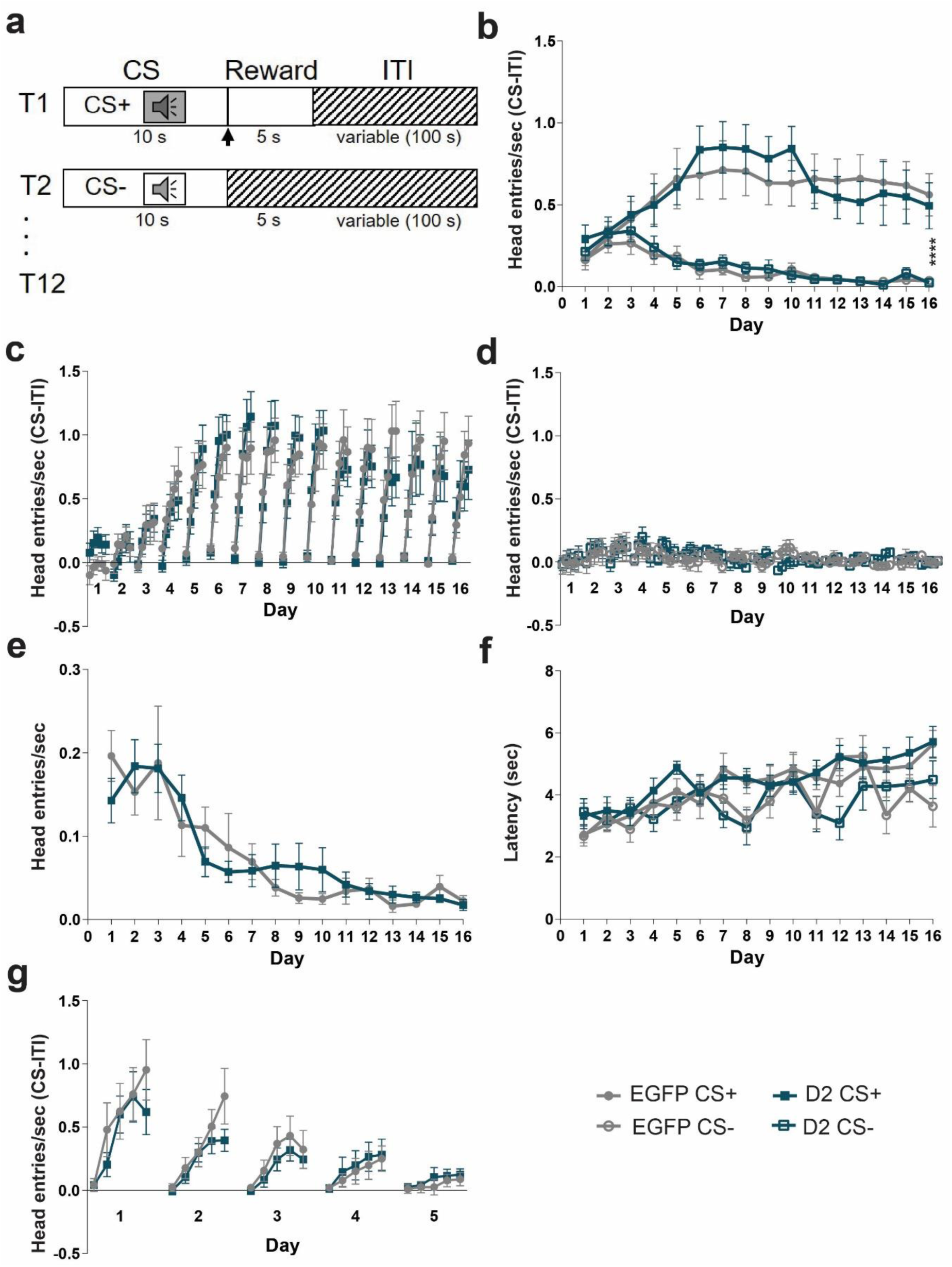
D2R overexpression does not impair Pavlovian learning. **(a)** Task design. Mice are trained with 24 (12 CS+, 12 CS-) trials /day for 16 days. Each trial starts with a 10 sec tone (CS+ or CS-). At the end of the CS+ a dipper comes up presenting milk as a food reward for 5 sec. There is an intertrial interval (ITI) variable in length (100 sec). **(b)** To determine learning, anticipatory responses (head entries/sec (CS-ITI)) were measured. Anticipatory head entries during the CS+ increased for both EGFP_NAcInd_ (filled light circles) and D2R-OE_NAcInd_ (filled dark squares) mice and decreased during the CS-for both EGFP_NAcInd_ (open light circles) and D2R-OE_NAcInd_ (open dark squares) mice. Both groups learned to distinguish the CS+ and CS- (\*\*\*\**p*< 0.0001). There was no effect of genotype on learning over the 16 days of training (*p*= 0.9977). **(c)** Within each day, the 10 sec CS+ was split into 2 second time bins to better visualize the anticipatory response. EGFP_NAcInd_ (filled light circles) and D2R-OE_NAcInd_ (filled dark squares) sharply increased head entries into the reward port across the 10 sec CS+ (*p*< 0.0001) and continued this anticipatory behavior across the 16 days of training (*p*< 0.0001). There was no effect of genotype on learning (*p*= 0.9375). **(d)** EGFP_NAcInd_ (open light circles) and D2R-OE_NAcInd_ (open dark squares) showed no anticipatory response during the 10 sec CS-. There was no effect of genotype on learning (*p*= 0.9988). **(e)** Head entries during the ITI decreased over the 16 days of training comparably for both groups (*p*= 0.6173). **(f)** Latency for the first head entry during the CS+ and CS- increased for EGFP_NAcInd_ (filled light circles/ open light circles) and D2R-OE_NAcInd_ (filled dark squares/ open dark squares). There was no difference between groups. **(g)** Anticipatory response to the CS+ attenuated over 5 days of extinction learning. Both groups similarly decreased anticipatory head entries during the CS+ (*p*< 0.0001). There was no difference in extinction learning between the two groups (*p*= 0.5247). Data from 16 animals per genotype was used to calculate all statistics reported.

To better visualize the pattern of anticipatory head entries we plotted head entries during the CS+ and CS- in 2 second bins. As mice learn the fixed duration of the CS+, their anticipatory response sharply increased during the 10 second CS+ (**Figure 3c**). We found that both D2R-OE_NAcInd_ and EGFP_NAcInd_ mice comparably increased their head entries over the duration of the CS+ (RM ANOVA: F(1, 15)= 470.5, *p*< 0.0001) and continued this pattern of anticipatory behavior across the 16 days of training (RM ANOVA: F(15,15)= 4.565, *p*<0.0001). However, there was no interaction between days, head entries during the CS+ and viral manipulation (**Figure 3c**, RM 3-way ANOVA: F(15, 15)= 0.5077, *p*=0.9375). Similarly, we observed no effect of day, head entries during CS- presentation and viral manipulation (**Figure 3d**, RM 3-way ANOVA: F(15, 15)= 0.2368, *p*= 0.9988). In addition, we observed no effect of viral manipulation (D2R vs EGFP) on the rate of head entries during the variable intertrial interval (ITI) (**Figure 3e**, RM ANOVA: F(15, 450)= 0.8535, *p*=0.6173). To further quantify their ability to learn the length of both cues, we measured the latency of first head entry with cue onset. For both groups, this latency increased only during the CS+ over the 16 days of conditioning, but no difference between groups (**Figure 3f**). This result further indicates that the mice have learned both the cue that predicts the reward and the timing of the reward that follows.

To test the effects of D2R upregulation on eliminating a conditioned response, we ran our 2^nd^ cohort of mice through 5 days of Pavlovian extinction. To reinstate the conditioned behavior, the mice received 3 days of Pavlovian conditioning as previously described (**Figure 3a**). For extinction, the protocol was identical to Pavlovian conditioning except now the mice no longer received a reward following the CS+. Both D2R-OE_NAcInd_ and EGFP_NAcInd_ mice attenuated their anticipatory head poking during the CS+ each day with near complete extinction by Day 5 (**Figure 3g**). Both groups had a comparable decay in their anticipatory head entries during the 10 second CS+ over the 5 days of extinction testing (RM 2-way ANOVA: F(4,4)= 6.536, *p*<0.0001). However, there was no interaction between day, CS+ anticipatory head entries and viral manipulation (RM 3-way ANOVA: F(4,4)= 0.8083, *p*= 0.5247), suggesting that D2R upregulation has no effect on extinction learning.

entries during CS- presentation and viral manipulation (**Figure 3d**, RM 3-way ANOVA: F(15, 15)= 0.2368, *p*= 0.9988). In addition, we observed no effect of viral manipulation (D2R vs EGFP) on the rate of head entries during the variable intertrial interval (ITI) (**Figure 3e**, RM ANOVA: F(15, 450)= 0.8535, *p*=0.6173). To further quantify their ability to learn the length of both cues, we measured the latency of first head entry with cue onset. For both groups, this latency increased only during the CS+ over the 16 days of conditioning, but no difference between groups (**Figure 3f**). This result further indicates that the mice have learned both the cue that predicts the reward and the timing of the reward that follows.

To test the effects of D2R upregulation on eliminating a conditioned response, we ran our 2^nd^ cohort of mice through 5 days of Pavlovian extinction. To reinstate the conditioned behavior, the mice received 3 days of Pavlovian conditioning as previously described (**Figure 3a**). For extinction, the protocol was identical to Pavlovian conditioning except now the mice no longer received a reward following the CS+. Both D2R-OE_NAcInd_ and EGFP_NAcInd_ mice attenuated their anticipatory head poking during the CS+ each day with near complete extinction by Day 5 (**Figure 3g**). Both groups had a comparable decay in their anticipatory head entries during the 10 second CS+ over the 5 days of extinction testing (RM 2-way ANOVA: F(4,4)= 6.536, *p*<0.0001). However, there was no interaction between day, CS+ anticipatory head entries and viral manipulation (RM 3-way ANOVA: F(4,4)= 0.8083, *p*= 0.5247), suggesting that D2R upregulation has no effect on extinction learning.

Next, to determine if D2R-OE_NAcInd_ mice have issues learning to withhold responses, we first trained mice to press a lever to earn a food reward and then tested them in a Go/No-Go task. Mice learned to press a lever to receive a milk reward using a continuous reinforcement (CRF) paradigm. Over the two days of training, both D2R-OE_NAcInd_ and EGFP_NAcInd_ mice were able to complete all trials (30 or 60) during the allotted time (60 min) (**Figure 4a)**. However, both groups improved their performance from Day 1 to Day 2. The number of successful trials (% Rewarded) increased (**Figure 4b**) for both the EGFP_NAcInd_ and D2R-OE_NAcInd_ mice (RM ANOVA: F(1,14)= 22.12, *p*= 0.0003) while there was no group effect of the viral manipulation (RM ANOVA: F(1, 14)= 0.2216, *p*= 0.6451). Both EGFP_NAcInd_ and D2R-OE_NAcInd_ mice got faster at the task as measured by a decrease in lever press latency (**Figure 4c**, RM ANOVA: F(1,14)= 15.76, *p*= 0.0014) with no difference between groups (RM ANOVA: F(1, 14)= 0.2255, *p*= 0.6422). We conclude that D2R upregulation has no effect on learning to press a lever for a food reward as we observed no difference in performance between the two groups.

**Figure 4.**
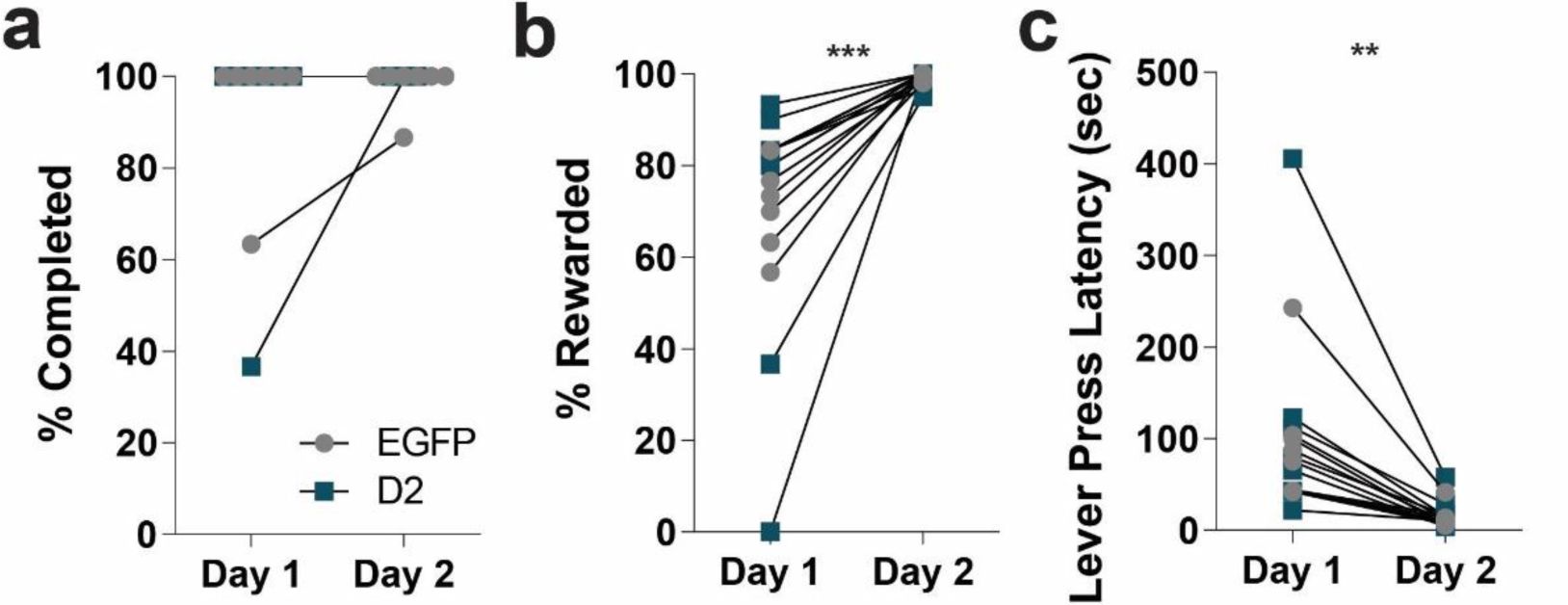
D2R upregulation does not affect reinforcement learning. Mice were trained to press a lever to obtain a milk reward on a continuous reinforcement (CRF) schedule for two days. **(a)** EGFP_NAcInd_ (light circles) and D2R-OE_NAcInd_ (dark squares) were able to complete all CRF trials on both days of training. **(b)** The number of rewarded trials (% Rewarded) comparably increased for EGFP_NAcInd_ (light circles) and D2R-OE_NAcInd_ (dark squares) mice over the 2 days of training (*** *p*= 0.0003). There was no difference between groups (*p*= 0.6451). **(c)** Lever press latency decreased from Day 1 to Day 2 for both groups (** *p*= 0.0014). There was no difference between groups (*p*= 0.6422). Data from 8 animals per genotype was used to calculate all statistics reported.

During the Go/No-Go task, mice use distinct visual cues to learn to either press a lever (“Go” trial) or withhold pressing of the same lever (“No-Go” trial) to receive a milk reward. In both trials, the lever is available for 5 seconds in which the animal must decide to press or not. A schematic of the task is shown in **Figure 5a**. The mice were first trained exclusively on Go trials and learning was established once they reached a criterion of 75% accuracy over three consecutive days. Next, No-Go trials were randomly intermixed with Go trials (60 total trials). Surprisingly, D2R-OE_NAcInd_ mice showed a deficit in the Go component of the task compared to the EGFP_NAcInd_ mice (**Figure 5b**, RM ANOVA: F(15, 210)= 2.029, p=0.0148). During the Go/No-Go task, both D2R-OE_NAcInd_ and EGFP_NAcInd_ control mice improved on withholding lever pressing during No-Go trials as measured by a decrease in correct responses (% incorrect) over the 35 days of testing (**Figure 5c**, RM ANOVA: F(34, 476)= 20.66, p<0.0001). However, there was no significant difference between the two groups (RM ANOVA: F (34, 476)= 0.4606, *p*=0.9965). Both groups also improved their performance on Go trials as measured by an increase in correct responses (% correct) the 35 days of testing (**Figure 5d**, RM ANOVA: F (34, 476)= 2.076, *p*=0.0005). Again, there was no difference in correct responses between the two groups over testing (RM ANOVA: F (34, 476)= 1.2, *p*=0.2073).

**Figure 5.**
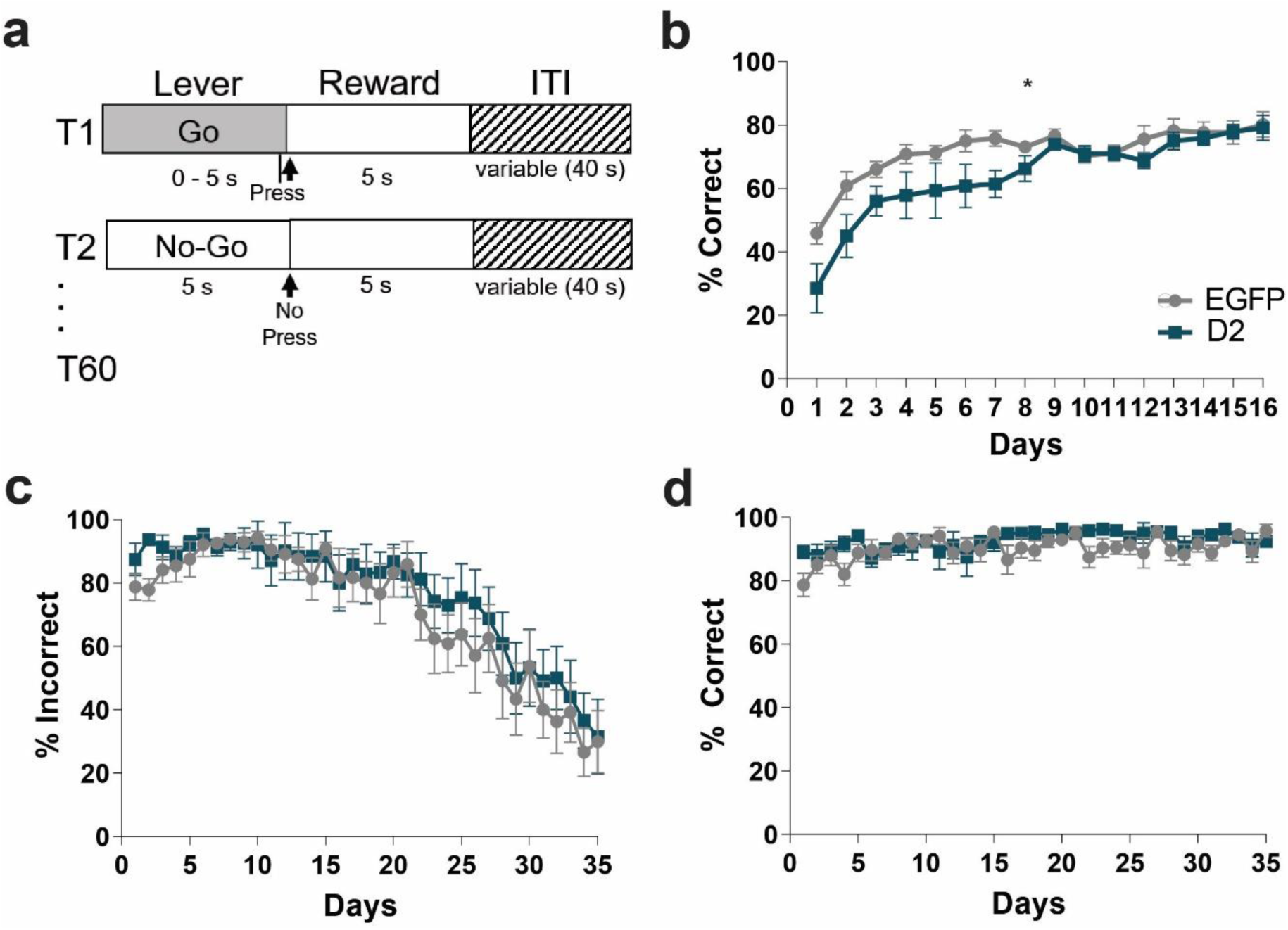
D2R upregulation does not impair No-Go learning. **(a)** Task design. Go trials: mice were trained to press a lever within 5 seconds to obtain a milk reward. No-Go trials: mice learned to withhold pressing the lever for 5 seconds to receive a milk reward. Mice only received a reward if they made a correct choice. Each session consisted of 30 Go and 30 No-Go trials that were randomly mixed. **(b)** Acquisition of Go trials only. D2R-OE_NAcInd_ (dark squares) showed a delay in the acquisition compared to EGFP_NAcInd_ (light circles) (* *p*= 0.0148). **(c)** Performance on No-Go trials during Go/No-Go testing. Both EGFP_NAcInd_ (light circles) and D2R-OE_NAcInd_ (dark squares) improved on No-Go trials and learned to withhold responding (decrease in % incorrect) over the 35 days of testing (*p<* 0.0001). There was no difference in performance between the two groups (*p*= 0.9965). **(d)** Performance on Go trials during Go/No-Go testing. Both groups performed better on Go trials (increase in % correct) over the 35 days of testing (*p*=0.0005). There was no difference in performance between groups (*p*= 0.2073). Data from 8 animals per genotype was used to calculate all statistics reported.

To determine whether D2R upregulation in this two cohorts leads to hyperlocomotion as has been described before (Donthamsetti et al., 2018; Gallo et al., 2018) we tested mice in the open field. D2R-OE_NAcInd_ mice showed an increase in locomotor activity in a standard 90-minute open field session replicating previous findings consistent with functional upregulation of D2Rs in iSPNs (**Figure 6**). D2R-OE_NAcInd_ mice continued to traverse the enclosure during the entire session while EGFP_NAcInd_ control mice attenuated their locomotion (**Figure 6a**, RM ANOVA: F(17, 510)= 5.231, *p*<0.0001). Furthermore, D2R-OE_NAcInd_ mice traveled a significantly greater total distance compared to controls (**Figure 6b**, *t*=2.705, *p*=0.0112). These results confirmed that our viral manipulation of over-expressing D2Rs in ventral iSPNs was functional.

**Figure 6.**
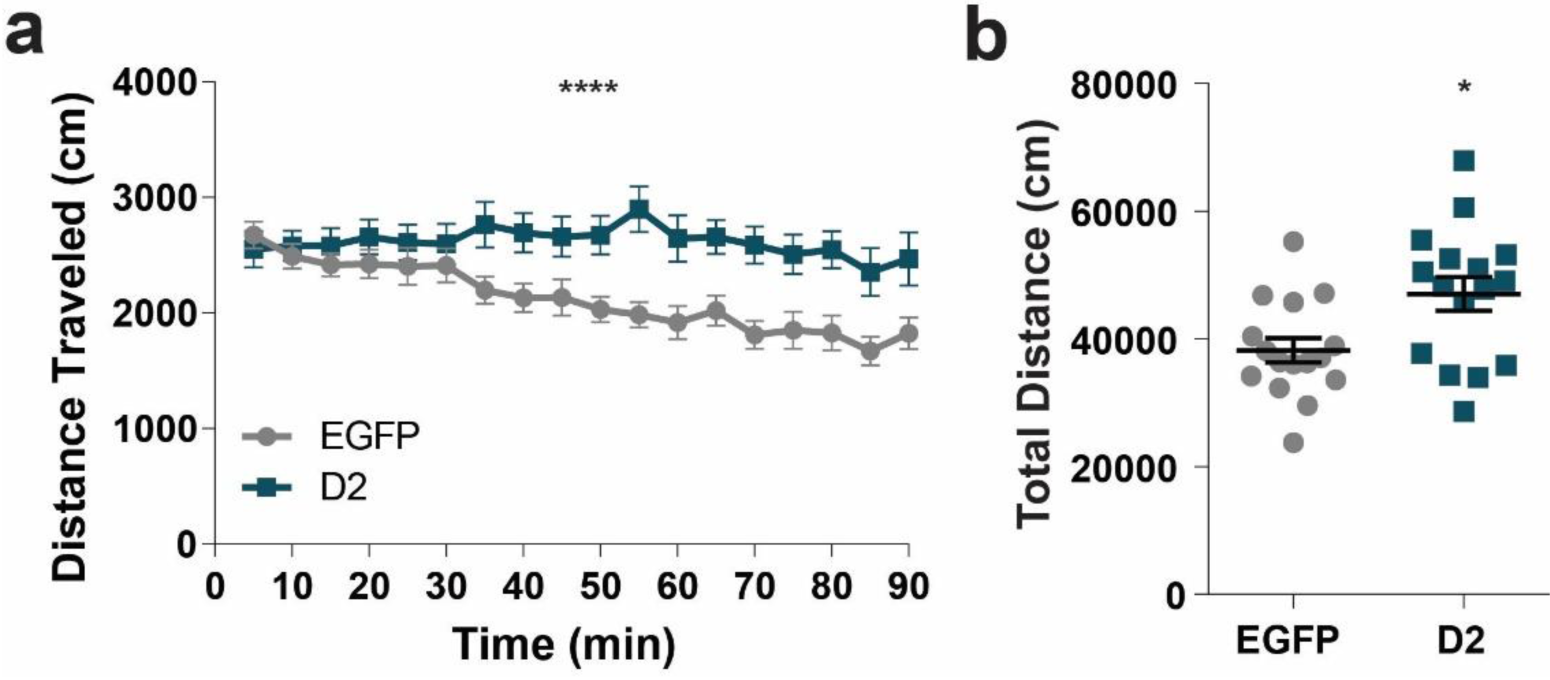
D2R upregulation induces hyperlocomotion. **(a)** D2R-OE_NAcInd_ (dark squares) mice traveled more distance in 5 min bins during a 90 min period than EGFP_NAcInd_ (light circles) mice (*p*< 0.0001). **(b)** D2R-OE_NAcInd_ mice traveled a greater total distance compared to EGFP mice (*p*= 0.0112). Data from 16 animals per genotype was used to calculate all statistics reported.

## Discussion

Here we determined if enhanced D2R expression in iSPNs of the NAc and the resulting deficit in iSPN synaptic transmission is important for associative reward learning and cognitive function in mice. To determine this, we used a viral approach to selectively over-express D2Rs in iSPNs of the NAc core and tested mice in an auditory Pavlovian conditioning task followed by a Go/No-Go paradigm. We found that upregulation of D2Rs in iSPNs in the NAc does not impair associative reward learning nor did it affect the extinction of response behavior when the Pavlovian cue was not paired with a reward any longer. In contrast to our initial hypothesis, the decreased function of the indirect pathway did also not cause deficits in No-Go learning, however, we did observe a slight delay in the acquisition of Go learning. Lastly, we replicated previous findings that D2R upregulation in NAc iSPNs enhances locomotion. These data suggest that while D2R upregulation in iSPNs enhances locomotor activity and the motivation to work for food (Donthamsetti et al., 2018; Gallo et al., 2018) it does not affect reward learning, at least under the conditions tested.

### D2R upregulation in NAc iSPNs does not impair Pavlovian conditioning and extinction learning

Accumulating evidence suggests that dopamine in the NAc core is important for associative reward learning. First, dopamine is released in response to reward predicting cues during a Pavlovian conditioning task (M. R. Bailey et al., 2018; Collins, Aitken, Greenfield, Ostlund, & Wassum, 2016). SPNs single-unit recordings from rats further showed that 75% of NAc neurons change their activity in response to reward-predicting cues. Of these neurons, half showed an increase in firing and half a decrease indicating a possible cell-type specific regulation consistent with D1R activation enhancing dSPN activity and D2R activation inhibiting iSPN activity (Day, Wheeler, Roitman, & Carelli, 2006). However, ventral striatal iSPN are not always inhibited after reward predicting cues. Pathway specific Ca^2+^ imaging revealed that reward-predicting cues enhanced iSPN activity in the lateral part of the ventral striatum, whereas it decreased iSPN activity in the ventro-medial striatum (Tsutsui-Kimura et al., 2017). This suggests that the regulation of iSPN and dSPN activity in response to cue induced dopamine is more complicated than the dichotomous model would suggest.

Second, inhibition of NAc core projecting DA neurons disrupts Pavlovian reward learning (Heymann et al., 2020). Contrasting this systemic administration of a D2R antagonist was found to enhance approach behavior and promoted learning in rats during an auditory Pavlovian conditioning paradigm (Eyny & Horvitz, 2003). However, the latter finding is difficult to interpret due to the systemic actions of the antagonist. Local infusion of dopamine receptor antagonists into the NAc revealed that D1R antagonism impaired memory consolidation during appetitive Pavlovian learning, whereas D2R antagonism had no effect on learning (Dalley et al., 2005). Similarly, local NAc core infusion of a D1R antagonist have been shown to impair Pavlovian Instrumental Transfer, whereas the infusion of a D2R antagonist had only mild effects (Lex & Hauber, 2008).

Third, animals demonstrate a divergence in approach behavior during Pavlovian learning that may be directed either towards the CS itself (sign-tracking) or the location of the reward delivery (goal-tracking). Dopamine is differentially involved in these approach behaviors; it is necessary for the development and expression of sign-tracking behaviors but only required for the expression of goal-tracking behaviors (Flagel et al., 2011). Furthermore, dopamine D1 and D2 receptors (D1Rs and D2Rs) play a differential role in approach behaviors. Both D2Rs and D1Rs play an important role in sign-tracking behaviors while only the activity of D1Rs is necessary for the development of goal-tracking behaviors (Roughley & Killcross, 2019). Here, our behavioral paradigm limited our analysis to only measure goal-tracking behaviors, which may explain why D2R upregulation did not affect Pavlovian learning. Taken together, these results implicate a role for NAc dopamine in associative reward learning using Pavlovian cues, however, the underlying mechanism seems to involve D1Rs rather than D2Rs. In so far, our results that D2R upregulation in iSPNs does not affect Pavlovian reward learning is not surprising.

Following the Pavlovian task, we tested D2R_NAcInd_ mice in extinction learning. We rationalized that reward omission during extinction learning should result in a dip in dopamine release in accordance with the reward predicting error model (Bromberg-Martin, Matsumoto, & Hikosaka, 2010). This dip in dopamine should decrease D2R mediated inhibition of iSPNs, thereby facilitating the learning of avoiding the Pavlovian response (Hikida, Kimura, Wada, Funabiki, & Nakanishi, 2010; Kravitz, Tye, & Kreitzer, 2012). Higher levels of D2Rs may make iSPNs more responsive to this regulation if under wild-type conditions all receptors are occupied by dopamine at baseline, whereas in the condition of D2R upregulation additional receptors are occupied. However, we found that that both D2R-OE_NAcInd_ and EGFP_NAcInd_ control mice attenuated their anticipatory head poking during the presentation of the previously reward predicting cue. Thus, D2R upregulation in iSPNs neither enhances nor impaired extinction learning.

### D2R upregulation in NAc iSPNs does not affect “No-Go” Learning

D2R-expressing iSPNs have been proposed to suppress movement or action initiation by gating the output of the basal ganglia via connections through the globus pallidus external (GPe) (Albin, Young, & Penney, 1989). Furthermore, inhibition of these neurons using the G_i_-coupled designer receptor hM4D_Gi_ enhances response initiation in a progressive hold own task, where the mouse as to hold down a lever for an increasing amount of time in order to obtain a food reward (M.R. Bailey et al., 2015; Carvalho Poyraz et al., 2016). Thus, we hypothesized that upregulation of G_i_-coupled D2Rs impairs learning if actions need to be suppressed. However, our Go/No-Go data did not reveal any impairment in No-Go learning and performance. During the Go/No-Go paradigm, both D2R-OE_NAcInd_ and control EGFP_NAcInd_ mice improved their performance and plateaued at the same level, demonstrating their ability to withhold responding during No-Go trials. This result shows that D2R upregulation in iSPNs is not sufficient to disrupt learning when actions need to be withheld.

Prior to Go/No-Go testing, the mice were trained exclusively on Go trials. During 16 days of Go training, the D2R-OE_NAcInd_ mice showed a mild impairment during the first 9 days. We believe that this impairment was due to a newly implemented time restriction for lever availability during Go training. During the preceding two days of continuous reinforcement (CRF) training, the lever was extended until the animal pressed the lever. However, during Go training the lever was only available for 5 seconds, and if no press was made the lever would retract and a new trial would start. Previous work from our group showed that D2R-OE_NAcInd_ mice are hyperactive in the open field (Gallo et al., 2015; Welch et al., 2019), a finding we replicated here. One possibility is that increased exploration in the operant chamber distracts D2R-OE_NAcInd_ mice from lever responding within the 5 sec time limit. Alternatively, D2R-OE_NAcInd_ mice have a deficit in recognizing and adapting to the changing circumstance of the 5 sec time limit.

### D2R upregulation in NAc iSPNs induces hyperlocomotion

D2R upregulation in NAc iSPNs increases locomotion (Gallo et al., 2015; Welch et al., 2019). To confirm that our manipulation was functional, we tested activity of D2R-OE_NAcInd_ and EGFP_NAcInd_ control mice in a standard open field box. We replicated hyperactivity in D2R-OE_NAcInd_ mice. Furthermore, post-hoc immunohistochemistry staining confirmed that our viral expression was targeted to the NAc core. Taken together, these results give us confidence that D2R upregulation was functional as in previous publication where we established with slice and *in vivo* electrophysiological measures that D2R upregulation impairs synaptic transmission of iSPNs collaterals in within the NAc and the canonical projections to the ventral pallidum.

We set out to determine how altered D2R levels affects associative and Go/No-Go learning in mice. We found that selective D2R upregulation on iSPNs in the NAc core does not impair associative reward learning and No-Go learning. In our previous published studies, we found that D2R upregulation enhances the willing ness to work for food if response efforts are high (Donthamsetti et al., 2018; Gallo et al., 2018). In contrast, both the Pavlovian conditioning and the Go/No-Go task do not require much effort to perform the task. Together with our previous published findings our data suggest that NAc D2Rs play a specific role in regulating motivation by balancing cost/benefit computations but does not affect associative learning (M. R. Bailey, Chun, Schipani, Balsam, & Simpson, 2020; Gallo et al., 2018; Simpson & Kellendonk, 2017; Ward et al., 2012).

## Acknowledgements

This work has been supported by RO1MH093672 (C.K) and RO1MH093672S1 (K.M.).

## Notes

### Competing Interest Statement

The authors have declared no competing interest.

